# First record of monogenean fish parasites in the Upper Lufira basin (Democratic Republic of Congo): dactylogyrids and gyrodactylids infecting *Oreochromis mweruensis, Coptodon rendalli* and *Serranochromis macrocephalus* (Teleostei: Cichlidae)

**DOI:** 10.1101/2022.06.15.496317

**Authors:** Gyrhaiss Kapepula Kasembele, Auguste Chocha Manda, Emmanuel Abwe, Antoine Pariselle, Fidel Muterezi Bukinga, Tine Huyse, Michiel W.P. Jorissen, Emmanuel J. Vreven, Wilmien J. Luus-Powell, Willem Smit, Joseph Roderick Sara, Jos Snoeks, Maarten P.M. Vanhove

**Affiliations:** Unité de Recherche en Biodiversité et Exploitation durable des Zones Humides (BEZHU), Faculté des Sciences Agronomiques, Université de Lubumbashi, Haut-Katanga, Democratic Republic of Congo; Department of Biology, Royal Museum for Central Africa, Leuvensesteenweg 13, BE-3080 Tervuren, Belgium; ISEM, Univ Montpellier, CNRS, IRD, Montpellier, France; Faculty of Sciences, Mohammed V University in Rabat, Rabat, Morocco; Section de Parasitologie, Département de Biologie, Centre de Recherche en Hydrobiologie, Uvira, Democratic Republic of Congo; Laboratory of Biodiversity and Evolutionary Genomics, Department of Biology, University of Leuven, Ch. Deberiotstraat 32, BE-3000 Leuven, Belgium; Research Group Zoology: Biodiversity & Toxicology, Centre for Environmental Sciences, Hasselt University, BE-3590 Diepenbeek, Belgium; DSI-NRF SARChI Chair (Ecosystem Health), Department of Biodiversity, University of Limpopo, Private Bag X1106, Sovenga, 0727, South Africa; Capacities for Biodiversity and Sustainable Development, Operational Directorate Natural Environment, Royal Belgian Institute of Natural Sciences, Vautierstraat 29, 1000 Brussels, Belgium; Department of Botany and Zoology, Faculty of Science, Masaryk University, Kotlářská 2, CZ- 611 37 Brno, Czech Republic; Zoology Unit, Finnish Museum of Natural History, University of Helsinki, P.O.Box 17, Helsinki FI-00014, Finland

**Keywords:** Lake Tshangalele, Haut-Katanga, *Cichlidogyrus*, *Enterogyrus*, *Gyrodactylus*, *Scutogyrus*

## Abstract

**Background:** Monogenean parasites have never been formally reported on fish from the Lufira basin. Then it is hypothesised that multiple monogenean species are to be recorded that are new to the region. This study aimed to record the gill monogenean parasite fauna of three cichlid fish species in the Upper Lufira basin by inventorying their diversity (species composition) and analysing their infection parameters (prevalence, mean intensity and abundance).

**Methods:** *Oreochromis mweruensis, Coptodon rendalli*, and *Serranochromis macrocephalus* were selected for the study, given their economic value and their abundance in the Upper Lufira basin. Monogeneans were isolated from the gills and stomach, mounted on glass slides with either Hoyer’s medium or ammonium picrate-glycerin for further identification under a stereomicroscope, based on morphological analysis of genital and haptoral hard parts. Indices of diversity and infections parameters were calculated.

**Results:** A total of thirteen gill monogenean parasite species (*Cichlidogyrus dossoui, C. halli, C. karibae, C. mbirizei, C. papernastrema, C. quaestio, C. sclerosus, C. tiberianus, C. tilapiae, C. zambezensis, Scutogyrus gravivaginus, S*. cf. *bailloni* and *Gyrodactylus nyanzae*) and one stomach monogenean (*Enterogyrus malmbergi*) were reported. A species richness of S= 10 for *O. mweruensis*, S= 6 for *C. rendalli* and S= 2 for *S. macrocephalus* were recorded. Five parasite species were reported to be common amongst *O. mweruensis* and *C. rendalli*. The most prevalent parasite species were *C. halli* (P= 80.9%) on *O. mweruensis, C. dossoui* (P= 92.9%) on *C. rendalli* and *C. karibae* and *C. zambezensis* (both of which P = 9.1%) on *S. macrocephalus* with a respective mean infection intensity of 7.9 on *O. mweruensis*, 9.8 on *C. rendalli* and 5 and 15, respectively, on *S. macrocephalus*. Results of this study reported new host ranges for five parasites species (*C. quaestio, S*. cf. *bailloni, E. malmbergi* on *O. mweruensis, C. halli* on *C. rendalli* and *C. karibae* on *S. macrocephalus*) as well as new geographical records for three of them (*S*. cf. *bailloni, E. malmbergi, C. karibae*).

**Conclusions:** This study highlights the richness of monogenean communities in the Upper Lufira basin and is a starting point for future helminthological studies, e.g. on the use of fish parasites as indicators of anthropogenic impacts.

## Background

Across the African continent, the Congo basin harbours the greatest species richness of fish [1-2]. The Congo basin covers 3,747 320 km^2^, and drains most of the Democratic Republic of Congo and parts of some of its bordering countries (Angola, Zambia, Tanzania, Burundi, Rwanda, Central African Republic and Republic of Congo) and a small part of Cameroon [3]. The Congo basin includes different types of habitats and is subdivided into sections: Upper Congo (called Lualaba), Middle Congo, and Lower Congo [2,4-5]. One of the major tributaries in the Upper Congo drainage is the Lufira River [6]. The Lufira River is subdivided into three sections: the Upper Lufira (from the source of the river to Lake Koni), the Middle Lufira (from downstream Lake Koni to the Kyubo Falls), and the Lower Lufira (from downstream the Kyubo Falls to the Kamalondo Depression, at the junction with the Lualaba River) [5,7]. In order to provide hydroelectric power, two successive dams were built in the Upper Lufira River; this created two artificial Lakes, Tshangalele (1930) and Koni (1949) [8-10]. Lake Tshangalele, located about 35 km east of the town of Likasi, holds a variety of fish, and it is also an UNESCO Man and the Biosphere Reserve, rich in birdlife [11-12]. In the Lufira River, most studies undertaken on biodiversity focused on vertebrates such as fish and birds [13-16]. Vast and speciose communities, which are often dominated by less sizeable animals such as flatworms or various parasite taxa, remain understudied, as is the case all over the world [17-18]. In view of the high biodiversity of potential host species in the tropics, it can be expected that parasitological surveys there would lead to the recording of many parasite species, including species new to science [19-20]. This study focuses on monogenean fish parasites due to their diversity, wide distribution, high host-specificity and single-host lifecycle, rendering them interesting models for studying the extent of parasite biodiversity and the underlying diversification mechanisms [21]. Monogeneans are common parasitic flatworms (Platyhelminthes) mostly infecting fish, and sporadically aquatic invertebrates, amphibians, reptiles and a single species of mammal (the hippopotamus) [22-27]. Infection sites of monogeneans on fish are typically gills, fins and/or skin [28], however they are also found rarely in the stomach, urinary bladder, intestine, oral or nasal cavity, eyes and heart [29-30]. Because of their one-host lifecycle and their close relationship with their host species, many monogeneans are specialists, infesting only a single host species (oioxenous specificity), though others are generalists, infesting two or more host species (stenoxenous specificity) [31-33]. Mendlová and Šimková [34] used a more extensive number of categories of host specificity on the basis of the phylogenetic relationships among (cichlid) host species. Parasites can be: (1) strict specialists when infecting only one host species; (2) intermediate specialists when infecting two or more congeneric host species; (3) intermediate generalists when infecting noncongeneric cichlid species belonging to the same tribe; and finally (4) generalists, when infecting noncongeneric cichlid species of at least two different tribes. African cichlids (taking also into account the Levant) are known to harbour monogenean parasites belonging to six genera: *Enterogyrus* Paperna, 1963; *Urogyrus* Bilong Bilong, Birgi & Euzet, 1994; *Onchobdella* Paperna, 1968; *Scutogyrus* Pariselle & Euzet, 1995; *Cichlidogyrus* Paperna, 1960 (Dactylogyridea) and *Gyrodactylus* von Nordmann, 1832 (Gyrodactylidea). The latter four are ectoparasitic genera, and among them, *Cichlidogyrus* is the most species-rich group with more than 138 nominal species described to date [35-37]. This study aims to record the monogenean parasite fauna of three cichlid fishes in the Upper Lufira basin; these parasites were never formally reported from this region. Objectives include: (i) inventorying the diversity of gill monogenean communities, and (ii) analyzing infection parameters of these monogenean parasites.

## Methods

### Study area

This study was conducted in the Upper Lufira basin (Figure 1), which is localized across the mining hinterland area in the west of the Haut-Katanga province (in the south of the former Katanga province). The climate is of type AW6 following the classification of Köppen [38], a rainy tropical climate with a rainy season extending from November to April [39]. Most precipitation falls from December to March [40]. Fishing is done essentially for *Coptodon rendalli* (Boulenger, 1896), *Oreochromis mweruensis* Trewavas, 1983, *Serranochromis macrocephalus* Boulenger, 1899, *Clarias gariepinus* (Burchell, 1822) and *Clarias ngamensis* (Castelnau, 1861) [12, 41]. Captured fish are intended for human consumption, for a small part by the local population around the Upper Lufira basin, and for most part in bigger towns such as Likasi and Lubumbashi.

**Figure 1:**
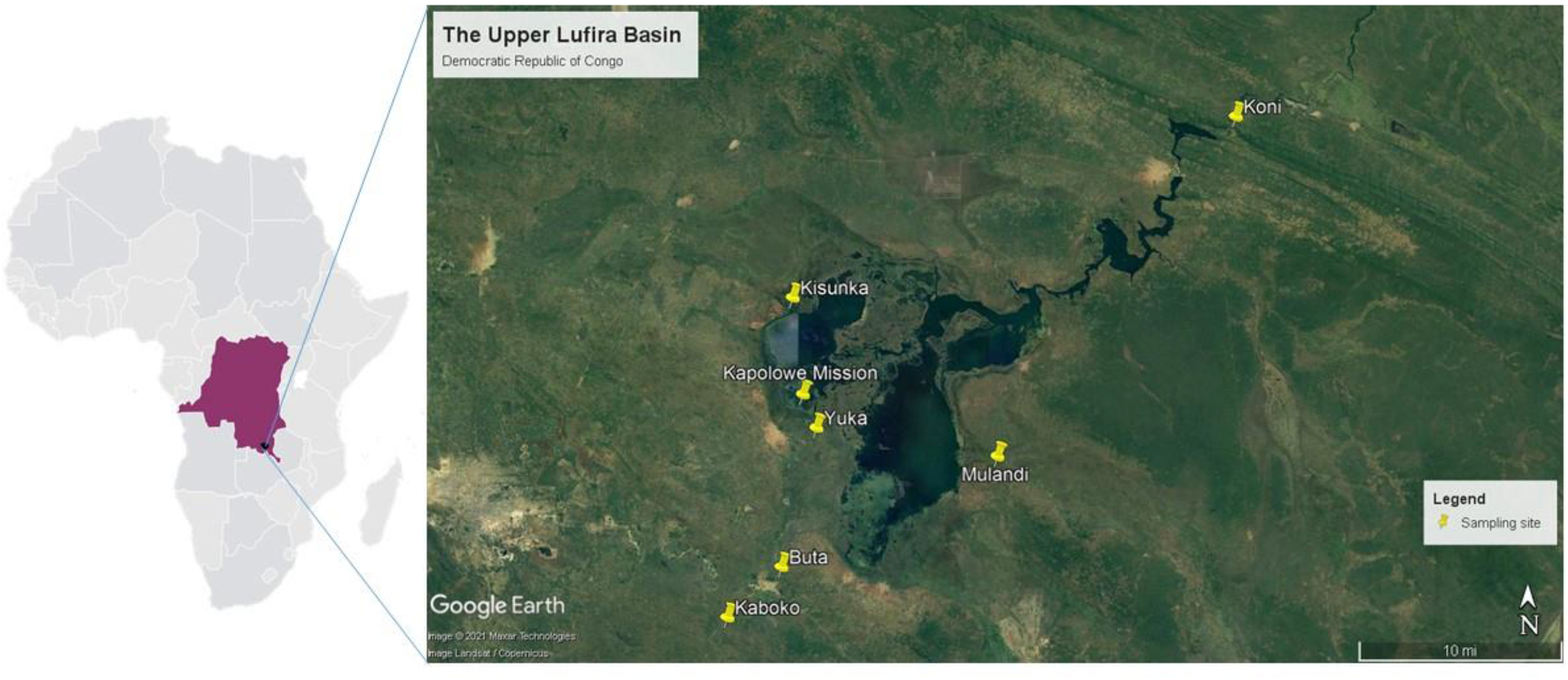
Map of sampling sites in the Upper Lufira basin: Lufira River (Kaboko 11°4’31.60”S; 26°55’2.40”E and Buta 11°2’21.60”S; 26°57’23.10”E); Lake Tshangalele (Kisunka 10°50’52.10”S; 26°57’50.60”E, Kapolowe Mission 10°54’59.50”S; 26°58’17.70”E, Yuka 10°56’25.30”S; 26°58’53.40”E and Mulandi 10°57’36.64”S; 27°6’44.88”E) and Lake Koni (Koni 10°43’3.65”S; 27°17’3.24”E)

### Fish sampling

Three fish species, *Oreochromis mweruensis, Coptodon rendalli* and *Serranochromis macrocephalus* were selected for the study, given their economic value and their abundance in the Upper Lufira basin [12, 41]. Fish were collected using nets or were bought from fishermen along the shores of the Lufira River, Lake Tshangalele and Lake Koni (Figure 1) between September 2015 and August 2018. Fish were kept alive in an aerated tank, and transported to a field laboratory. Fish were identified up to the species level following the keys by Skelton [42] and Lamboj [43]. Fish were killed by severing the spinal cord just posterior to the cranium, immediately prior to examination, following Olivier *et al*. [44]. Fish were processed as the total length (TL) and the standard length (SL) were measured to the nearest centimetre, and the weight was taken in gram for each fish.

### Parasite sampling

To collect monogenean parasites, fish were dissected and the right gill arches removed by dorsoventral section. One fish amongst all the fishes sampled was randomly dissected and inspected for monogenean parasites in its stomach. Gill arches and the stomach were placed in a Petri-dish containing water for examination using a stereomicroscope Optica 4.0.0. Parasites were dislodged from the gill filaments using entomological needles and fixed between a slide and cover slip into a drop of either Hoyer’s medium or ammonium picrate-glycerin (a preparation described by Malmberg, 1957) according to Nack *et al*. [45]. Twenty-four hours later, coverslips were sealed using nail varnish. Parasites were deposited in the invertebrate collection of the Royal Museum of Central Africa (RMCA) under accession numbers XXX.

### Monogenean community composition, indices of diversity and infection parameters

Morphological identifications of the retrieved parasite specimens were conducted based on the sclerotized parts of the haptor, the male copulatory organ (MCO) and the vagina, using an Optica BA310 and a phase-contrast Olympus BX50 microscope. Parasite identification up to species level, and comparison with known congeners was based on García-Vásquez *et al*. [46-47], Přikrylová *et al*. [48-49], Gillardin *et al*. [50], Muterezi *et al*. [51], Pariselle and Euzet [35,52], and Fannes *et al*. [53]. Parasite diversity was summarized by the species richness index (S), indices of Shannon (H) and Equitability of Pielou (J). Infection parameters: prevalence (P), mean intensity (MI) and mean abundance (MA) were provided following definitions given by Margolis *et al*. [54] and Bush *et al*. [55]. Statistical analysis was performed using Past 3.1 software.

## Results

Fish processed for the study had different size and weight range. For *Oreochromis mweruensis* (n=47) the mean TL was 18.2 ± 4.1 cm and 14.6 ± 3.2 cm for the mean SL, and the mean weight was 72.7 ± 38.8 g. For *Coptodon rendalli* (n = 28) the mean TL was 15.1 ± 2.8 cm and 12.0 ± 2.4 cm for the mean SL, and the mean weight = 72.7 ± 38.8 g. For *Serranochromis macrocephalus* (n = 11) the mean TL was 16.9 ± 3.4 cm and 14.0 ± 2.8 cm for the mean SL, and the mean weight was 81.9 ± 51.5 g.

### Monogenean community composition and indices of diversity in the Upper Lufira basin

Representatives of four genera of monogeneans, *Cichlidogyrus, Gyrodactylus* and *Scutogyrus* (on the gills) and *Enterogyrus* (in the stomach), were collected (Table 1). Among them were ten known species of *Cichlidogyrus*, one species of *Gyrodactylus*, two species of *Scutogyrus* and one species of *Enterogyrus*. Parasite diversity indices were reported to be 10, 6 and 2 for S; 1.5, 1.2 and 0.6 for H; and 0.6, 0.8 and 0.8 for J respectively for *O. mweruensis, C. rendalli* and *S. macrocephalus*. The distribution of monogeneans per sampling period or per season is shown in Table 2.

**Table 1:**
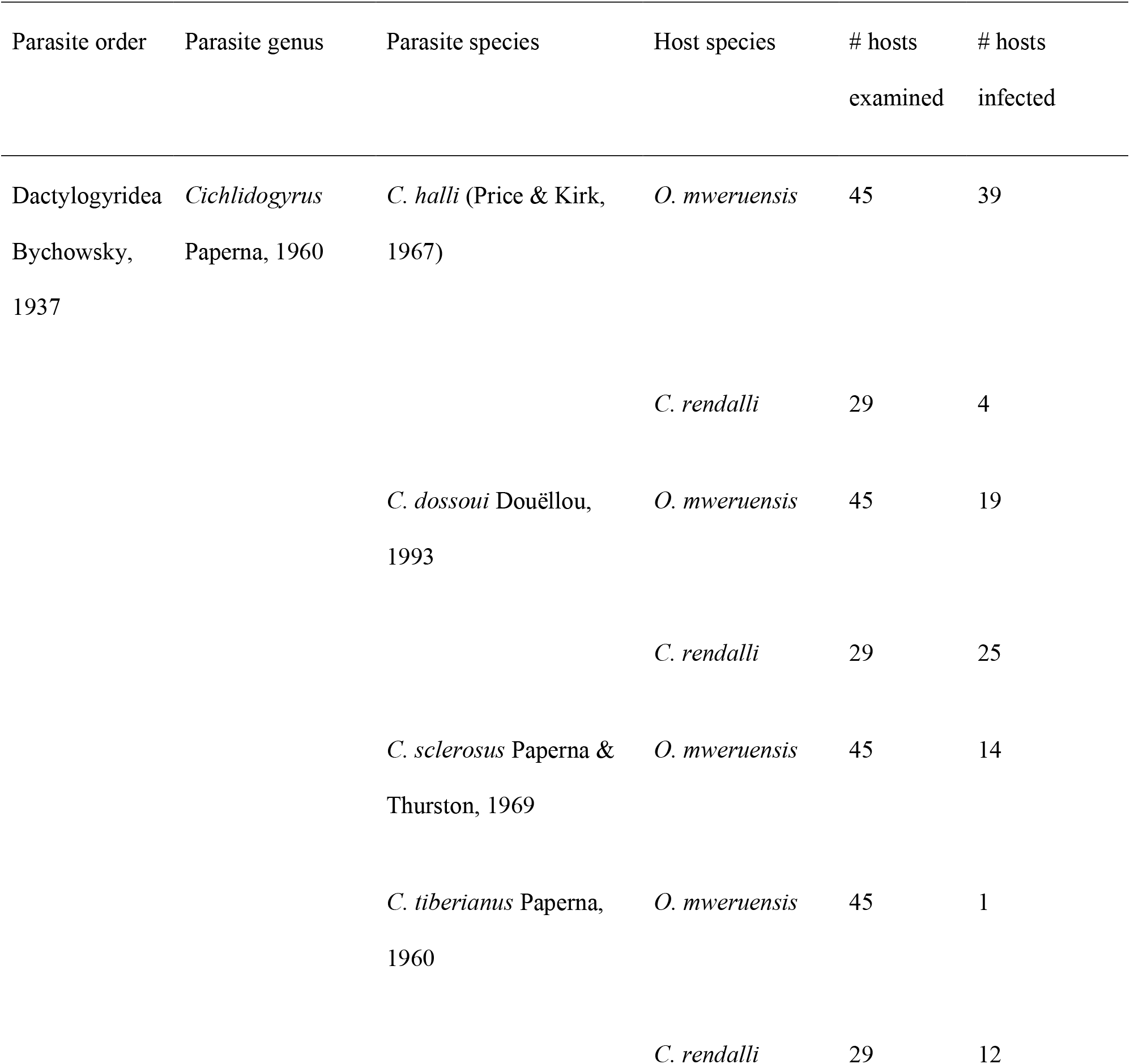
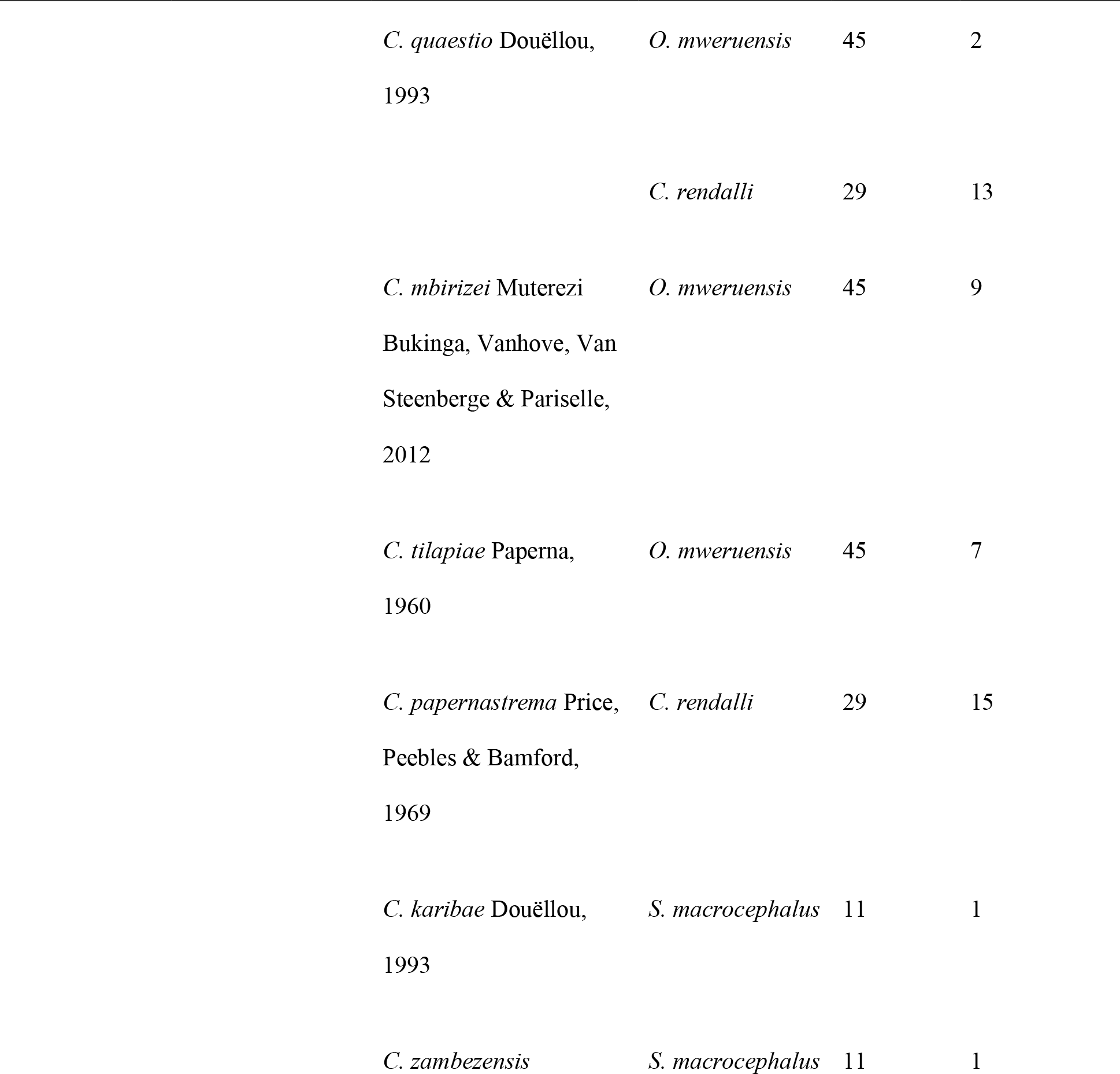
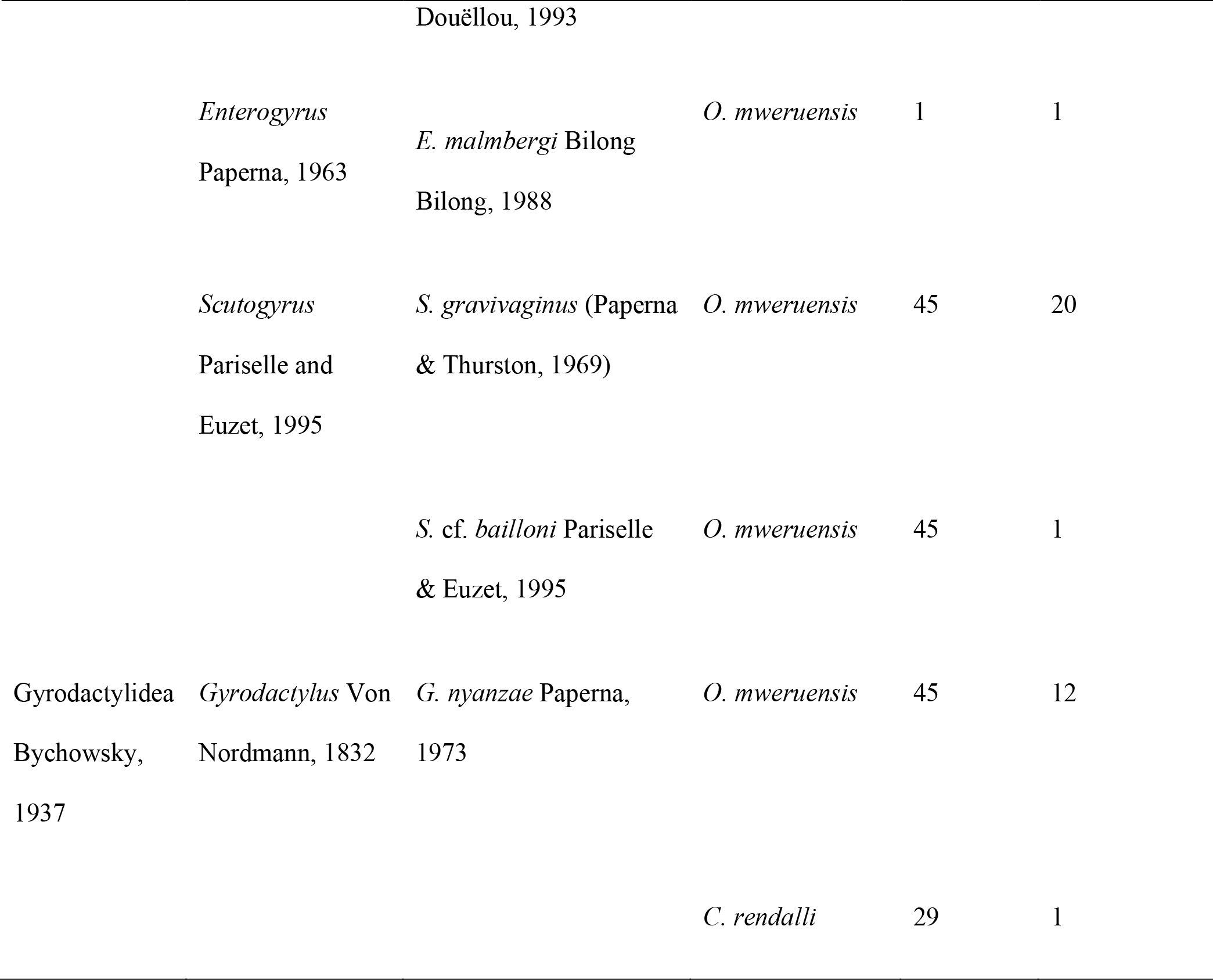
The monogenean parasite species recovered from *Oreochromis mweruensis, Coptodon rendalli* and *Serranochromis macrocephalus* in the Upper Lufira basin

**Table 2:**
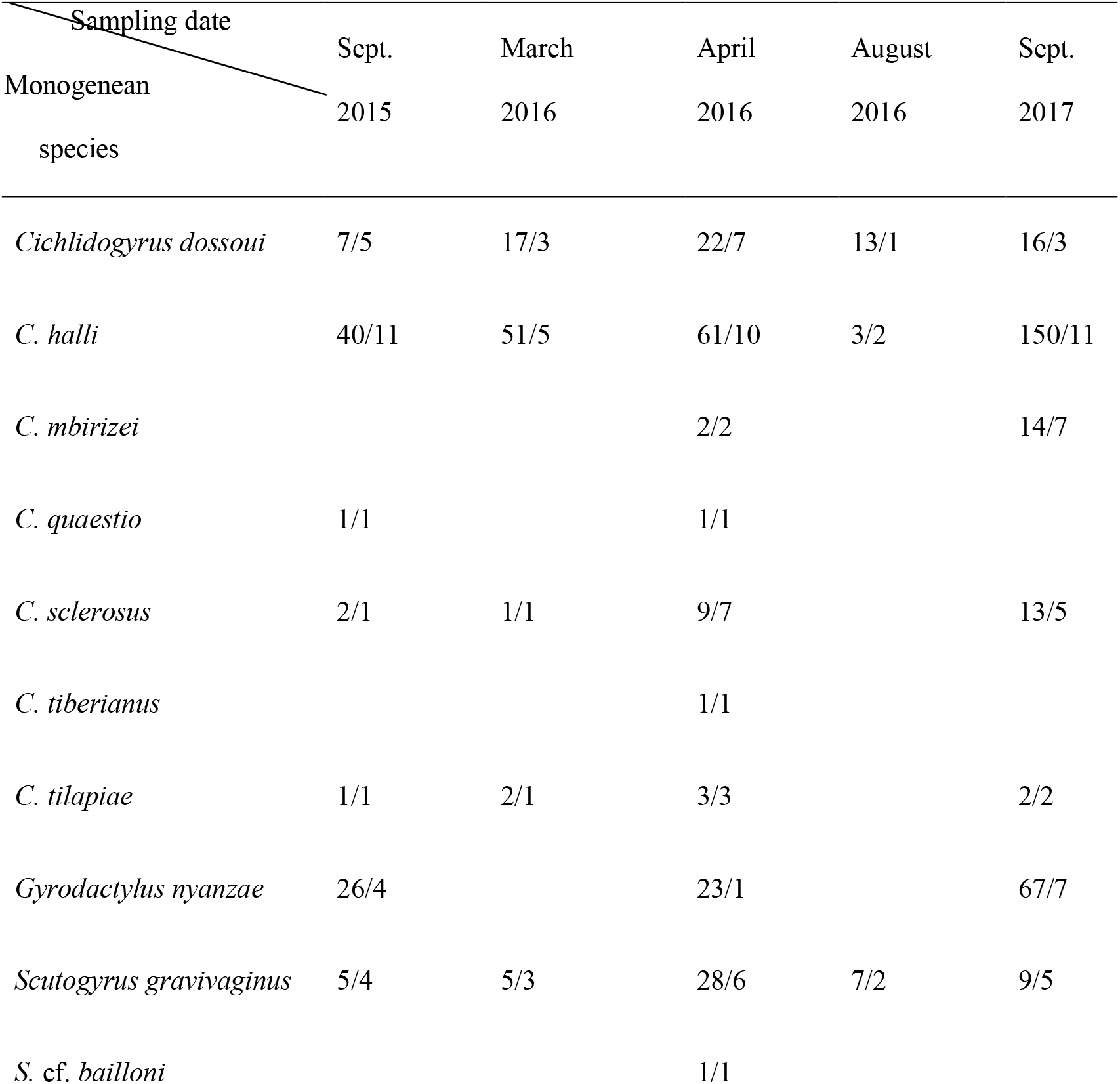
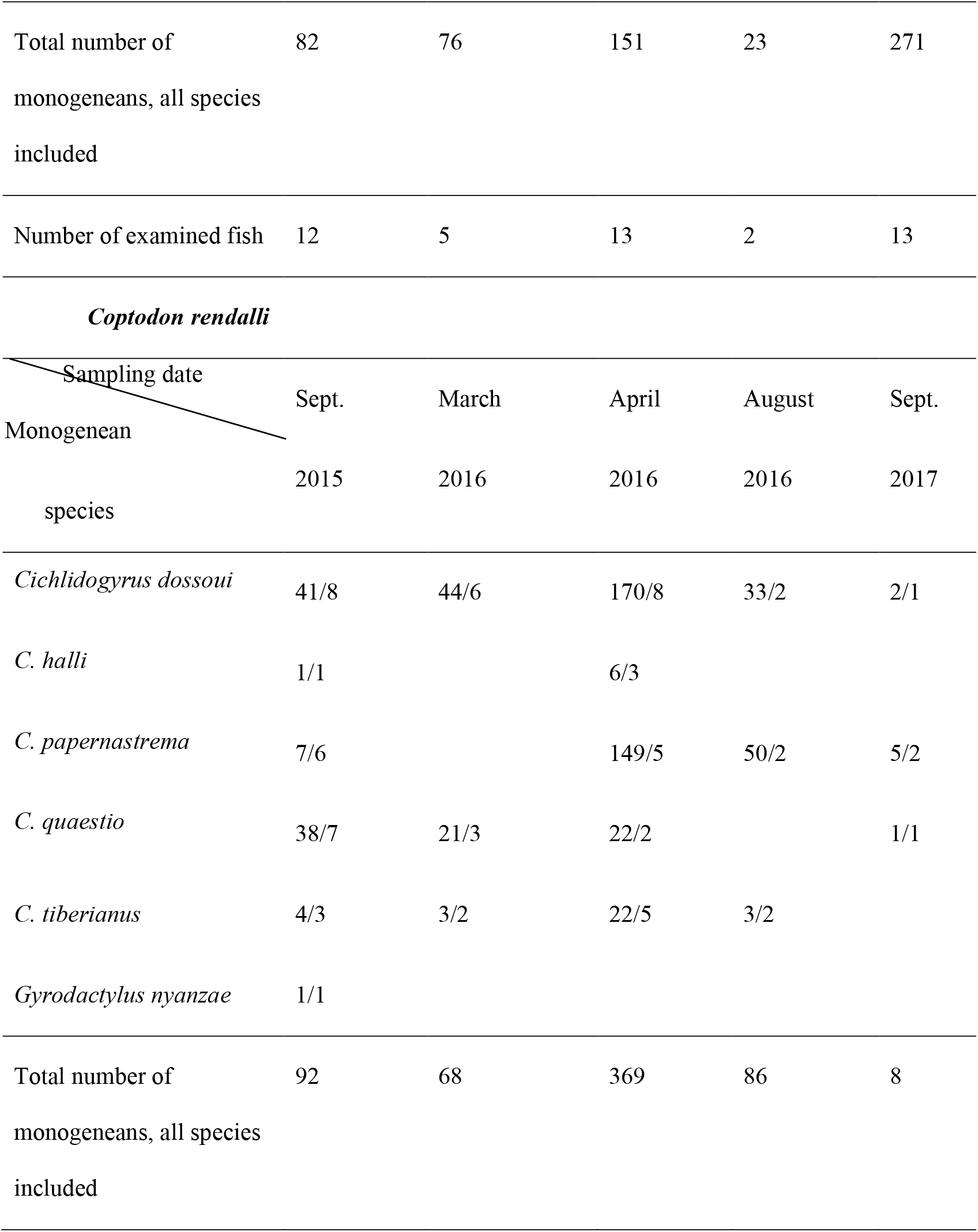
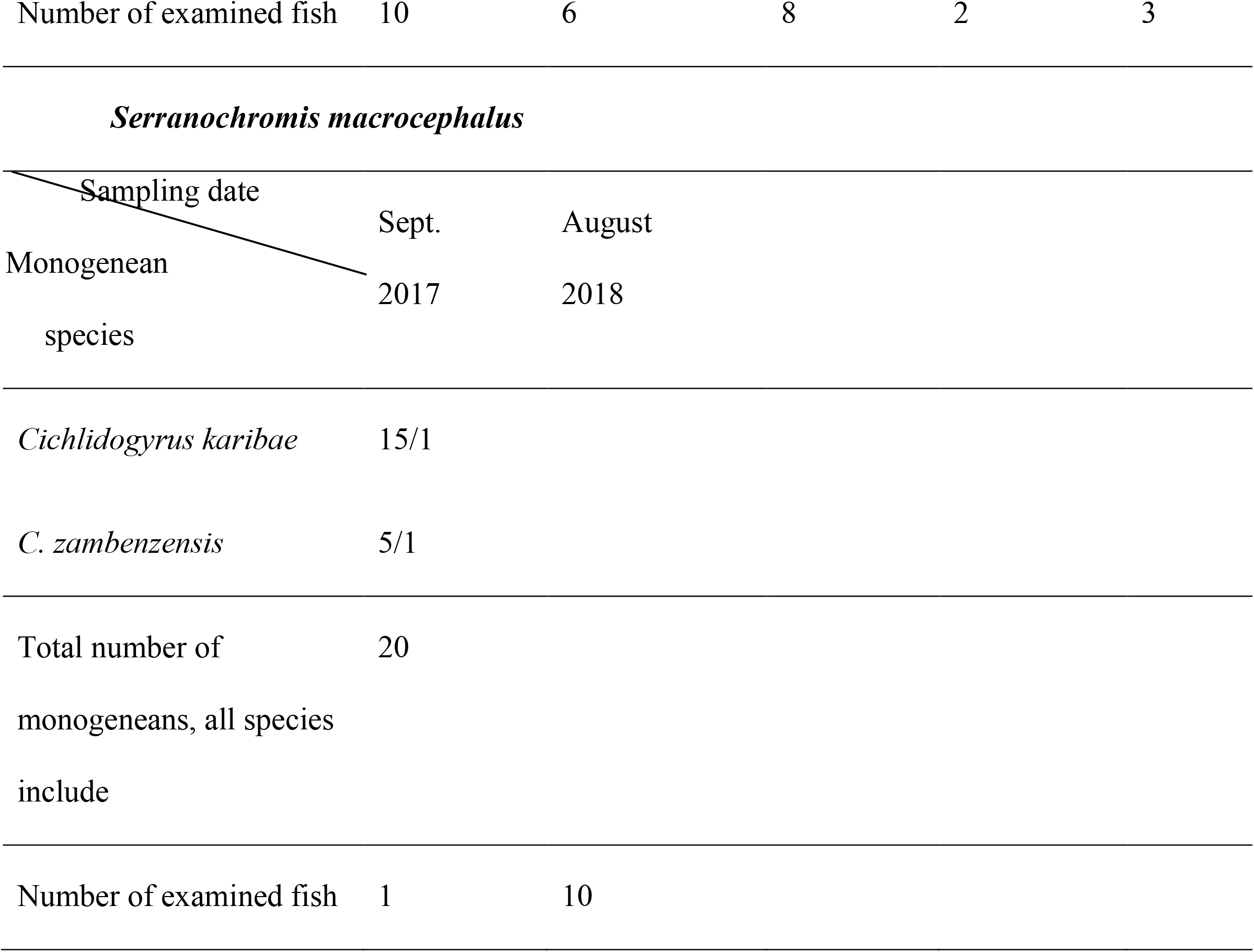
X/Y: Number of specimens of a given parasite species, out of the number of infected fish per host according to sampling period [August, September (Sept.): dry season; March, April: rainy season]

### Infection parameters of monogenean parasites in the Upper Lufira basin

Prevalence, mean intensity and mean abundance presented in this section take into account hosts grouped without seasonal subdivision.

The highest prevalences recorded was 80.9% for *C. halli* on *O. mweruensis*, 92.3% for *C. dossoui* on *C. rendalli*, and 9.1% for both *C. zambezensis* and *C. karibae* on *S. macrocephalus*. A low prevalence of 2.1% was recorded for *C. tiberianus, S*. cf. *bailloni* for *O. mweruensis*, and 3.8% for *G. nyanzae* from *C. rendalli* (Figure 2).

**Figure 2:**
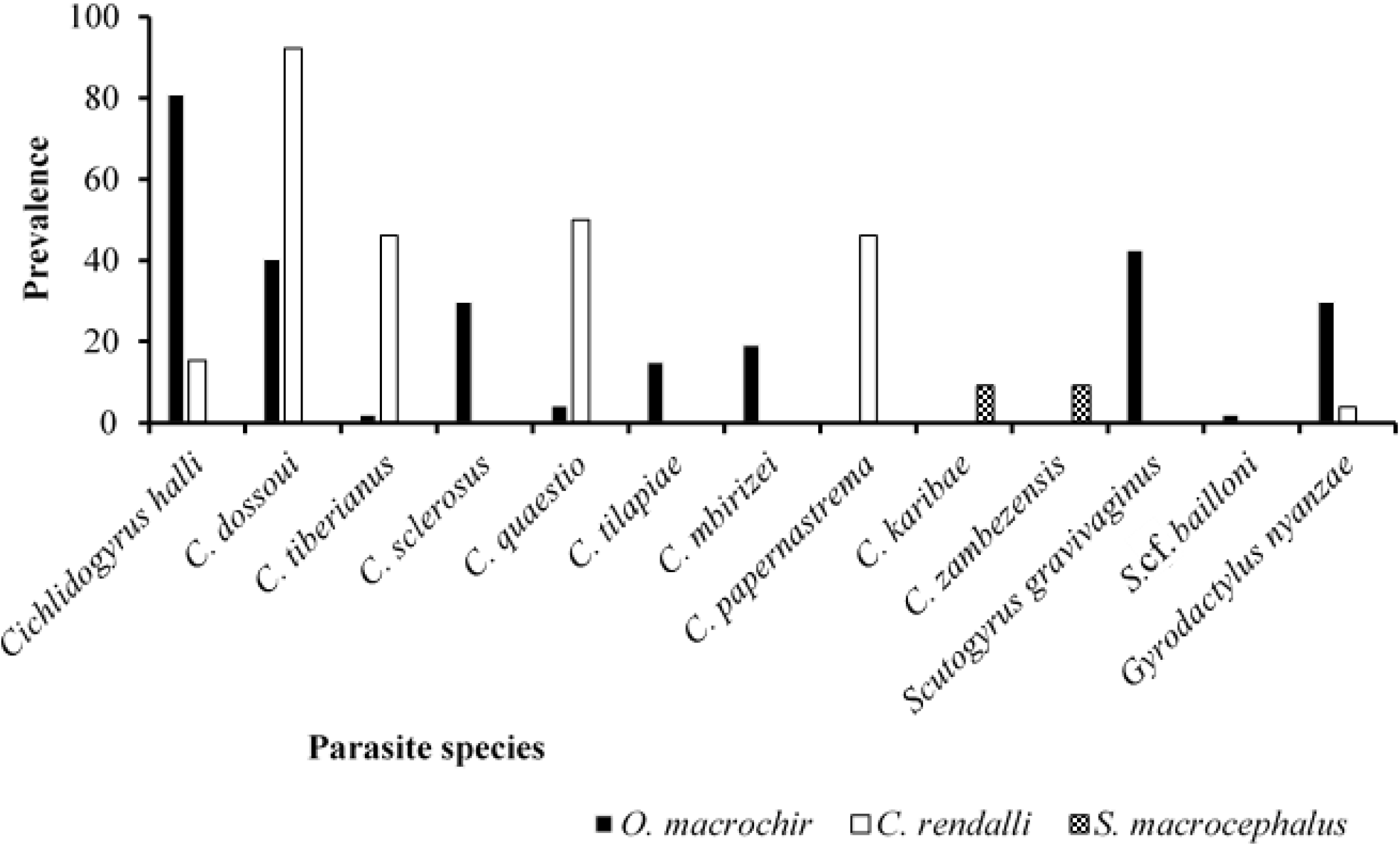
Parasite prevalence (%) per monogenean species recovered on the gills of *Oreochromis mweruensis, Coptodon rendalli* and *Serranochromis macrocephalus* in the Upper Lufira basin

For *G. nyanzae* the highest MI = 8.7± 9.9 was recorded from *O. mweruensis* and a low of MI= 1 ± 0 from *C. rendalli*. Conversely *C. papernastrema* obtained a MI of 17.1 ± 24 when examining the latter fish host. For *S. macrocephalus, C. karibae* was the parasite with the highest mean intensity (MI= 15) and *C. zambezensis* the lowest (MI= 5) (Figure 3).

**Figure 3:**
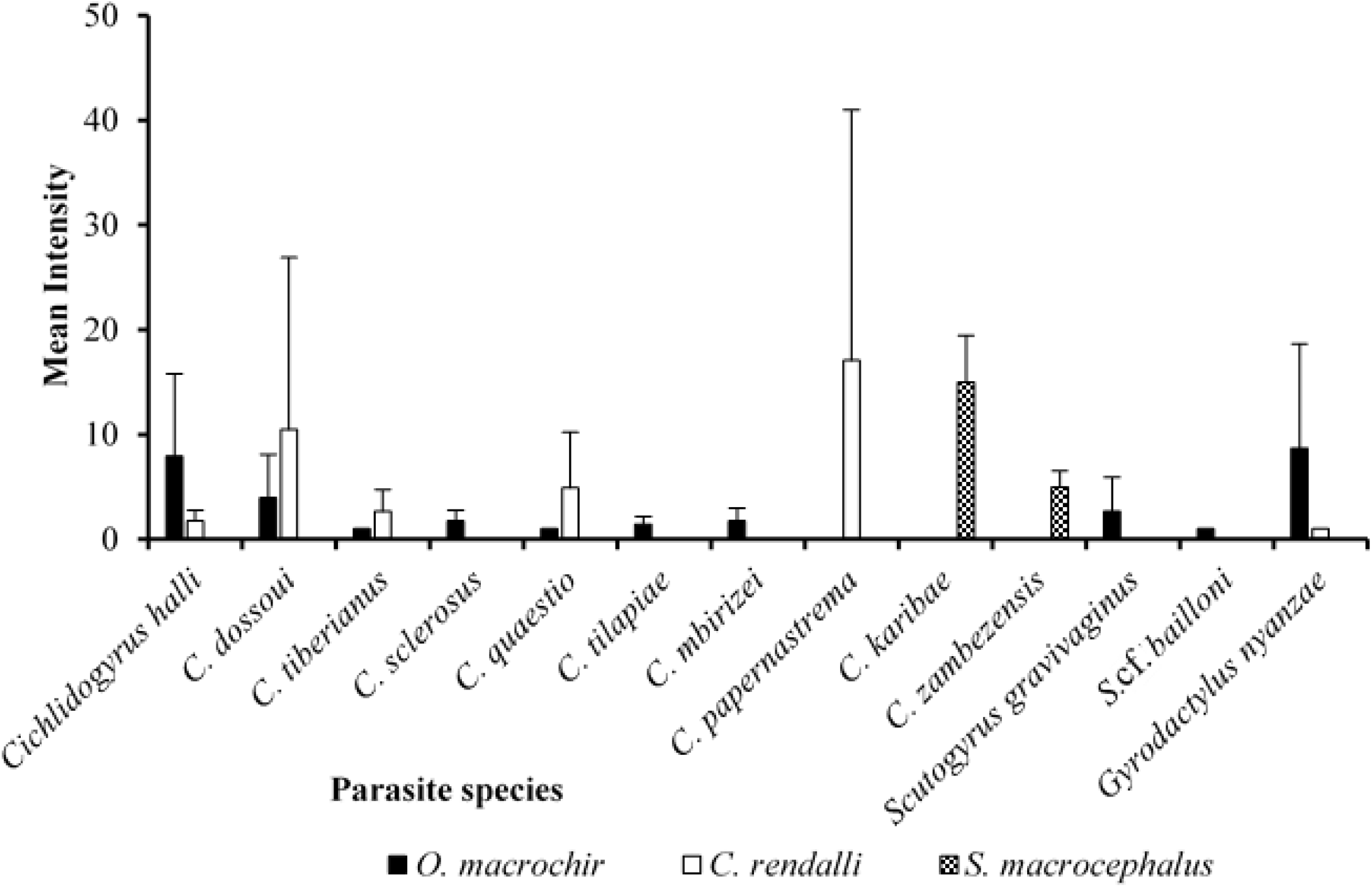
Mean intensity of each monogenean species recovered on the gills of *Oreochromis mweruensis, Coptodon rendalli* and *Serranochromis macrocephalus* in the Upper Lufira basin, with bars about the mean indicating the standard deviation

The results regarding the mean abundance reveal that on *O. mweruensis, C. halli* (MA= 6.4 ± 7.7) is the most abundant species; on the gills of *C. rendalli, C. dossoui* (9.7 ± 15.6) is the most abundant species; and the highest abundance of monogeneans on *S. macrocephalus* is 1.4 ± 4.5 per examined fish for *C. karibae* (Figure 4).

**Figure 4:**
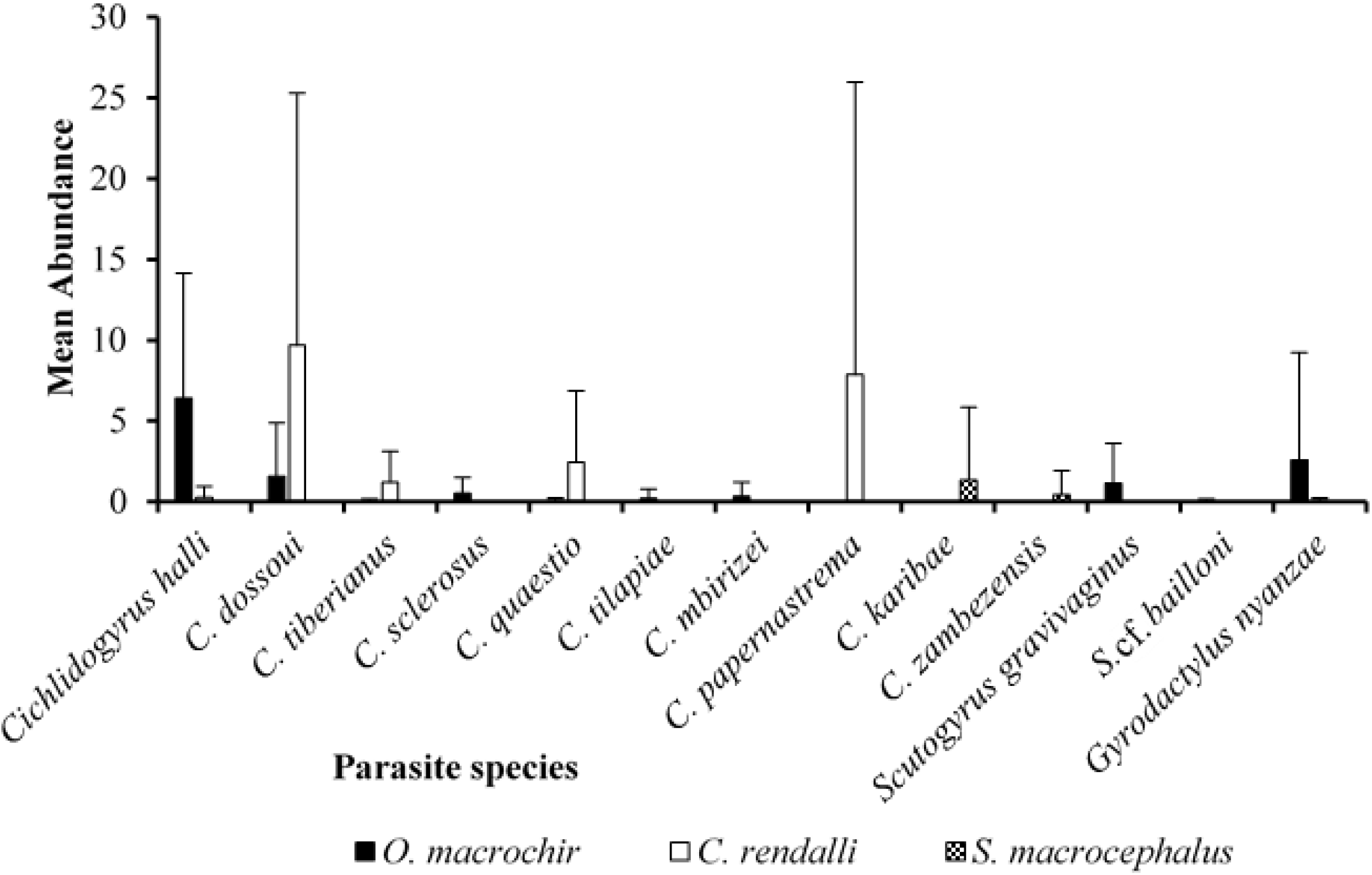
Mean abundance of each monogenean species recovered on the gills of *Oreochromis mweruensis, Coptodon rendalli* and *Serranochromis macrocephalus* in the Upper Lufira basin, with standard deviation

## Discussion

This research was conducted to explore the monogenean parasite fauna of three economically important and abundant cichlid species in the Upper Lufira basin, a part of the Upper Congo basin. In this study thirteen gill and one stomach monogenean species were recorded. Parasite species were already reported from fish belonging to the genera *Oreochromis, Coptodon* and *Serranochromis* [35,51, 56]. Although few studies on monogenean parasites from the Congo basin have been conducted in the Lake Tanganyika, Bangweulu-Mweru, Upper Lualaba, Kasai, Lower Congo and Pool Malebo Ecoregions (*sensu* Thieme *et al*. [57]) (e.g. Vanhove *et al*. [58]; Gillardin *et al*., [50]; Muterezi *et al*. [51]; Jorissen *et al*. [56, 59-60]; Geraerts *et al*. [61]), this study is the first to record monogenean parasites in the Lufira basin.

The known host range of five parasite species is extended in this study. *Cichlidogyrus quaestio, S*. cf. *bailloni* and *E. malmbergi* were recorded for the first time from *O. mweruensis*; *C. halli* from *C. rendalli*; and *C. karibae* from *S. macrocephalus. Cichlidogyrus karibae* was described by Douëllou [62] on *Sargochromis codringtonii* (Boulenger, 1908) in Lake Kariba (Zambezi basin, Zimbabwe). *Enterogyrus malmbergi* was described by Bilong Bilong [63] from the stomach of *Oreochromis niloticus* (Linnaeus, 1758) in the Sanaga River (Cameroon). *Scutogyrus bailloni* was formally described by Pariselle and Euzet [52] on *Sarotherodon galilaeus* (L, 1758) in the Mékrou River (Niger basin, Niger, West Africa). Since only a single similar parasite specimen was retrieved in this study on the gills of *O. mweruensis*, it cannot be assigned to *S. bailloni* with certainty. Nevertheless these (putative in case of *S. bailloni*) records substantially expand the known geographical distribution of these three monogenean species. Considering species richness, our results are similar to previous reports of monogenean gill parasites for these fishes in the Congo basin. In this study, ten monogenean species were found on *O. mweruensis*, while Jorissen *et al*. [56, 59] collected nine parasite species in the Bangweulu-Mweru ecoregion on *O. mweruensis* (of which seven are shared, except for *Cichlidogyrus mbirizei, C. quaestio* and *S*. cf. *bailloni* on *O. mweruensis* from the Lufira river system, and *C. cirratus* and *C. papernastrema* on *O. mweruensis* from the Bangweulu-Mweru ecoregion). Six monogenean species were found on *C. rendalli* in this study, while Jorissen *et al*. [59] collected five parasite species (all but *C. halli* corresponding to those found in this study) in the Bangweulu-Mweru ecoregion. On *S. macrocephalus*, two monogenean species (*C. karibae* and *C. zambezensis*) were found in this study while Jorissen *et al*. [59] reported only the last species, on fewer host fish.

In terms of infection parameters, on *O. mweruensis*, one parasite species had a prevalence higher than 50% in the Upper Lufira basin (*C. halli*, P= 80.9%) against two monogenean species in the Bangweulu-Mweru reported by Jorissen *et al*. [59] (P= 57.1% for *C. dossoui* and *S. gravivaginus*). On *C. rendalli, C. dossoui* (P= 92.3%) in the Upper Lufira basin, and *C. dossoui, C. quaestio* and *C. tiberianus* in the Bangweulu-Mweru, have P>50% following comparison with Jorissen *et al*. [59]. On *S. macrocephalus*, no parasite species had a prevalence higher than 50% in the Upper Lufira basin, while *C. zambezensis* reaches a prevalence of 100% in the Bangweulu-Mweru. Regarding the infection intensity (Table 1), on *O. mweruensis*, in the Upper Lufira basin, the most infected fish harbour up to 30 specimens of *C. halli*, followed by 25 specimens of *G. nyanzae*, against 37 parasite specimens of *G. nyanzae* and 21 parasite specimens of *C. cirratus* in Bangweulu-Mweru (reported by Jorissen *et al*. [59]). On *C. rendalli* in the Upper Lufira basin, the most infected fish harboured up to 84 specimens of *C. papernastrema*, followed by *C. dossoui* with 68 monogenean specimens against respectively 29 and 20 specimens of *C. dossoui* and *C. quaestio* in the Bangweulu-Mweru Ecoregion. Finally, on *S. macrocephalus* in the Upper Lufira, the most infected fish contain up to 15 and 5 parasite specimens of *C. karibae* and *C. zambezensis* respectively while Jorissen *et al*. [59] reported 21 parasite specimens of *C. zambezensis* in the Bangweulu Mweru. These differences in infection parameters may be due to sample size, season, biogeographical distribution or other environmental parameters, as communities of cichlid-infecting monogeneans have been observed to fluctuate e.g. seasonally and between habitat types, and parasite species composition may change between areas and basins [64-66].

## Conclusion

We reported stomach and gill monogenean species richness and infection parameters from three cichlid species in the Upper Lufira basin. A total of 13 monogenean species were recovered from *O. macrochir, C. rendalli* and *S. macrocephalus*. These findings are the first record of monogeneans in the Upper Lufira basin. For future sampling, it will also be interesting to study other groups of fish parasites other than monogenean parasites, as well as other fish species or families, to record the diversity of parasites [56, 59]. In addition, parasites can also be used as bioindicators of water quality [67-69] in this ecosystem where there is a substantial anthropogenic threat, especially from mine pollution [70-71]. The use of parasites as bioindicators of environmental conditions has been applied previously on African cichlids [72]. This study can serve as a baseline whereby future studies conducted on fish from the Upper Lufira basin can be compared to this study so as to establish if there has been a change in parasite composition and parasite load over time.

## Acknowledgements

VLIR-UOS is thanked for supporting this study through the South Initiative « *Renforcement des capacités locales pour une meilleure évaluation biologique des impacts miniers au* Katanga (RD Congo) *sur les poissons et leurs milieux aquatiques* », the local team of the University of Lubumbashi (BEZHU) namely C. Kalombo Kabalika, P. Kiwele Mutambala, B. Katemo Manda, M. Kasongo Ilunga Kayaba and C. Mukweze Mulelenu for their help in fish sampling, the international team, I. Přikrylová, for her contribution in confirmation of parasite identification, and F.A.M. Volckaert (KU Leuven) and L. Janssens de Bisthoven (Royal Belgian Institute of Natural Sciences) for hosting G.K. Kasembele in their teams during his respective research visits to Belgium. Finally, we thank the *Institut Congolais pour la Conservation de la Nature* (ICCN) for facilitating and authorising sampling.

## Availability of data and materials

Slides of monogenean parasites are available in the invertebrate collection of the Royal Museum of Central Africa, Tervuren, Belgium.

## Authors’ contributions

ACM, JS and MPMV designed and supervised this study. ACM, EA, EJV contributed to sampling, the collection and identification of fish. FMB, WJLP, WS, JRS and MPMV helped with the collection and preparation of the gill parasites. AP, MWPJ, MPMV helped with the morphological identification of parasites species. MPMV helped with the writing of the paper, analysis of the data, interpretation and discussion of results and provided scientific background in the field of monogenean research. All the authors critically read and edited the manuscript, and approved the final manuscript.

## Ethics approval and consent to participate

Fish were collected using nets or were bought from fishermen. In the absence of relevant animal welfare regulations in the DRC, we had used the guidelines and authorization in accordance with the Unité de Recherche en Biodiversité et Exploitation durable des Zones Humides (BEZHU) of the Université de Lubumbashi

## Consent for publication

Not applicable

